## Supplementary Information for "Structural Conservation of Insulin/IGF Signalling Axis at the Insulin Receptors Level in *Drosophila* and humans"

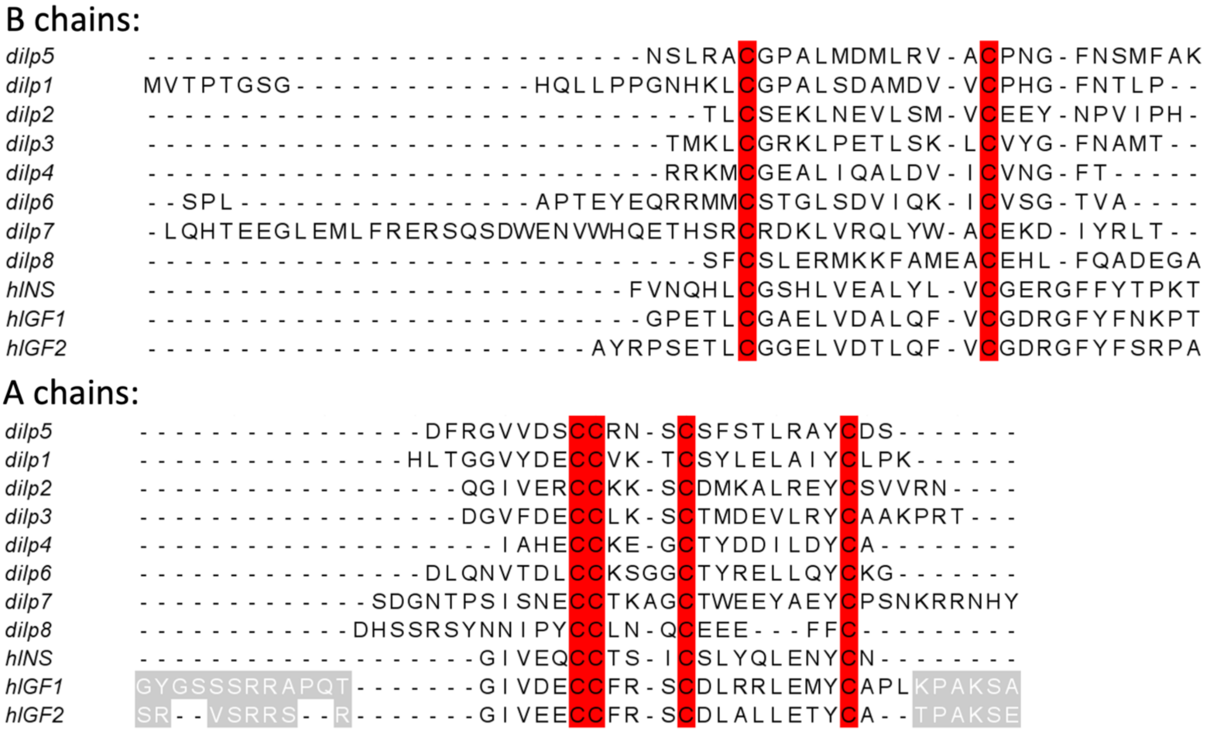

**Supplementary Information Figure 1. Sequence alignment of DILP1-8, human insulin (hINS) and human IGF1 (hIGF1) and IGF2 (hIGF2)**. C- and D-domains of IGFs are in gray (in A-chains panel).

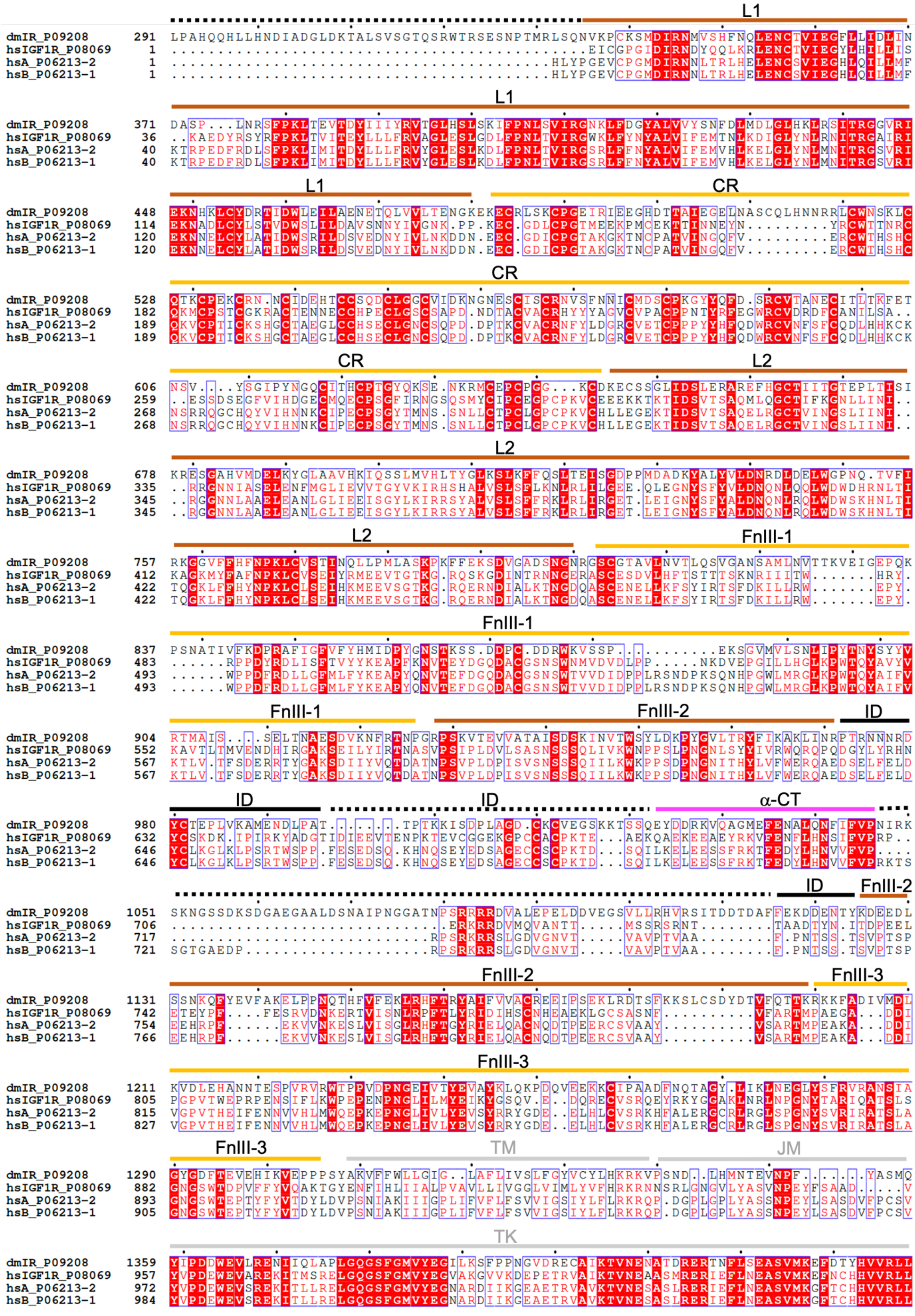
L1
CR
L2
FnIII-1
FnIII-2
ID
α-CT
FnIII-3
TM
JM
TK
dmIR_P09208
hsIGF1R_P08069
hsA_P06213-2
hsB_P06213-1

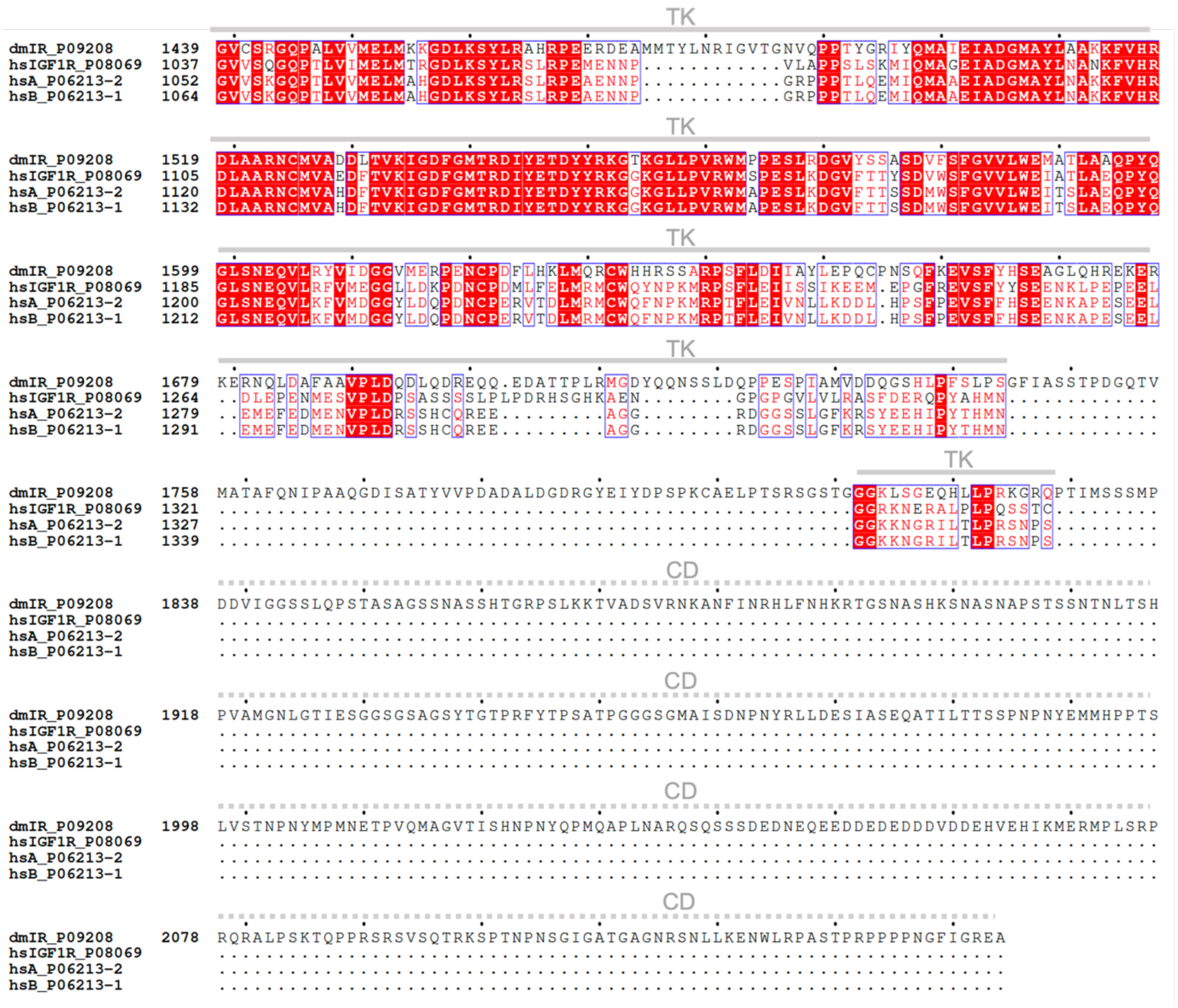

**Supplementary Information Figure 2. Sequence alignment of dmIR, hIR-A(hsA)/-B (hsB) isoforms, and hIGF-1R (hsIGF1R).** The dmIR-ECD studied here spans the 291-1309 sequence. Brown, yellow, magenta and black lines mark the span of the dmIR-ECD individual domains based on its cryoEM structure reported here; dotted black lines – regions not seen in the cryoEM map. Expected TM, JM and TK regions are indicated by grey lines, while dmIR-unique CD region is indicated by grey dotted lines.

**Supplementary Information Figure 3. CryoEM data processing workflow and structure solution.**

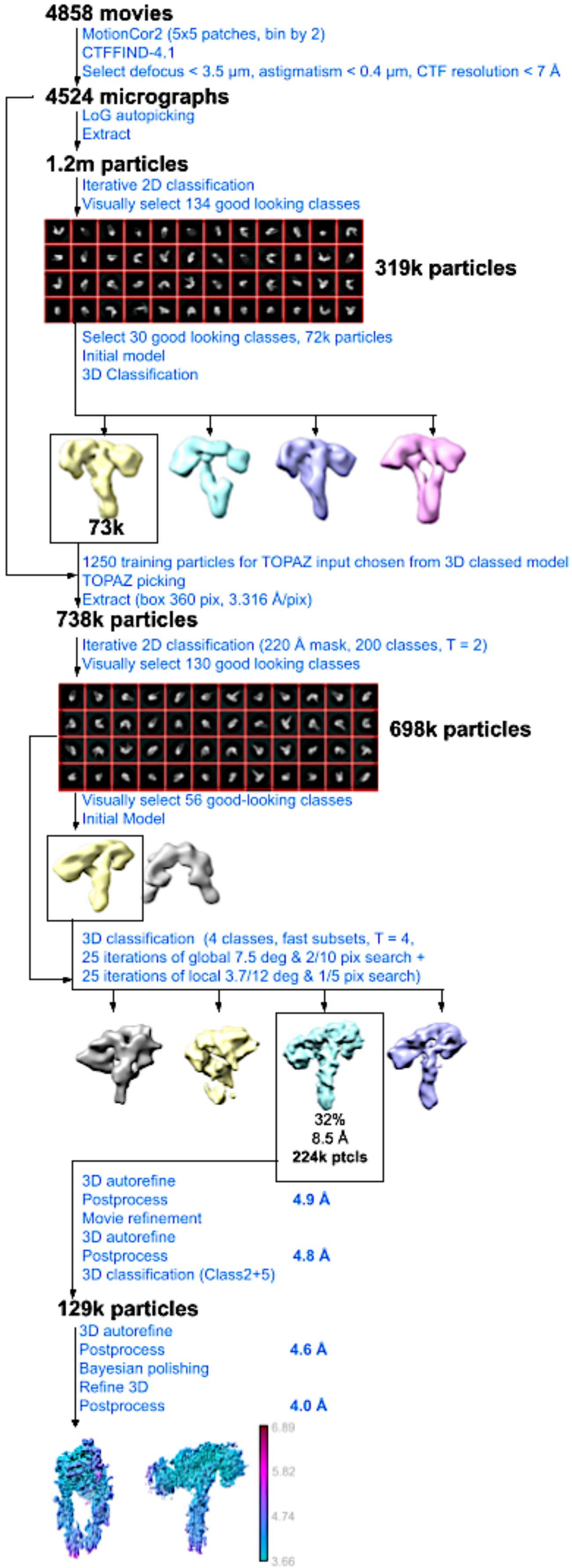

**Supplementary Information Note 1. RMS deviations of the superpositions of the dmIR-ECD and hIR corresponding domains for some representative hIR structures (PDB IDs are given), protomers and the whole ECDs.** All superpositions were carried out by the LSQ option in Coot[1] on the Cα atoms, except the global superpositions of the whole ECDs for which SSM option in Coot was used. The range of target Cα atoms is given for each type of superposition.

The 827-847 and 881-887 loops in dmIR-ECD were removed prior to the superposition to minimise their potentially misleading bias in the superposition of the 'core' folds of the selected FnIII-1 domains.

Approximate boundaries of the domains in dmIR-ECD and hIR:

| | **dmIR-ECD domains:** | **human IR-A domains:** |
|---|---|---|
| L1 | 335-480 | 1-154 |
| CR | 482-644 | 155-309 |
| L2 | 645-800 | 310-469 |
| FnIII-1 | 803-926 | 470-593 |
| FnIII-2 | 928-971 116-1200 | 593-635 758-807 |
| CT | 1021-1041* | 691-720 |
| FnIII-3 | 1193-1310 | 811-910 |

*region visible in the map

***(A) Dynamic protomer FnIII-1 domain superpositions.***

Targets: Ala807-Asn925 in dmIR (B-chain) on Glu471 Glu-Asp591 in hIR (mice IR for 7S* PDBs); loops 827-847 and 881-887 deleted for superposition of FnIII-1 and FnIII-1' domains to remove dmIR loops-specific bias. For the IGF-1R the target were Ala807–Asn925 (dmIR B-chain) and Ser461-Ile572 for the IGF-1R (PDB ID: 6PYH[2] and 7S0Q[3] (IGF-1R protomer).

| | | | |
|---|---|---|---|
| 6HN5[4] | 1.19 (F chain) | 7SL2[8] | 1.46 (A chain) |
| 6SOF[5] | 1.77 (A chain) | 7SL4[8] | 1.26 (A chain) |
| 6PXV[6] | 1.16 (A chain) | 7STJ[8] | 1.42 (A chain) |
| 7PG0[7] | 1.86 (B chain) | 7STK[8] | 1.52 (A chain) |
| 7PG2[7] | 2.95 (B chain) | 7STI[8] | 1.69 (A chain) |
| 7PG3[7] | 1.60 (B chain) | 7MQS[9] | 1.52 (E chain) |
| 7PG4[7] | 1.53 (B chain) | 6PYH[2] | 1.97 (A chain) |

***(B) Static protomer FnIII-1 domain superpositions.***

Targets: Ala807 - Asn925 in dmIR (A-chain) on Glu471 - Asp591 in hIR (mice IR for 7s* PDBs)

| | | | |
|---|---|---|---|
| 6SOF | 1.51 (C chain) | 7PG3 | 1.84 (A chain) |
| 6HN5 | 1.33 (E chain) | 7SL2 | 1.35 (B chain) |
| 7MQS | 1.30 (F chain) | 7SL4 | 1.60 (B chain) |
| 7PG0 | 1.66 (A chain) | 7SL7 | 1.19 (A chain) |
| 7PG2 | 1.90 (A chain) | 7STI | 1.60 (B chain) |
| 7PG4 | 1.90 (A chain) | 7STJ | 1.55 (B chain) |

7STK 1.33 (B chain)
6PYH 2.34 (D chain)
7S0Q 1.61 (A chain)

***(C) Site 1 L1 domains – on the lower arm on dynamic protomer.***
Targets: Pro338 – Thr476 in dmIR-ECD (B-chain) and Val7 – Val146 in hIR

7SL2 1.71 (A chain)
7SL4 1.05 (A chain)
7SL7 0.97 (A chain)
7STK 0.92 (A chain)
6SOF 1.39 (A chain)
6HN5 1.04 (E chain)
7MD4[10] 0.98 (B chain)

***(D) Site 1' L1 domains – on the upper arm on static protomer.***

6HN5 1.10 (E chain)
6SOF 1.45 (C chain)
7PG0 1.66 (A chain)
7PG2 1.72 (A chain)
7PG4 2.05 (A chain)
7PG3 1.55 (A chain)
7SL2 1.13 (B chain)
7SL4 1.24 (B chain)
7SL7 1.14 (A chain)
7STI 1.17 (B chain)
7STJ 1.20 (B chain)
7STK 1.25 (B chain)
7MQS 1.48 (E chain)
7S0Q 1.03 (A chain)
6PYH 1.05 (D chain)

***(E) SSM global superpositions of dmIR-ECD and representative hIR (mIR – 7s*) and hIGF-1R (6PYH).***
PDB ID, number of atoms in dmIR and in IR (IGF-1R) used in the alignment, respectively, and number of residues superimposed.

6HN5 4.64, 1853, 1653, 917
6SOF 5.06, 1853, 1924, 773
6PXV 5.36, 1853, 1830, 852
7PG0 5.27, 1853, 1742, 1016
7PG2 5.20, 1853, 1700, 1063
7PG3 4.34, 1852, 1721, 884
7PG4 4.62, 1853, 1677, 913
7SL2 4.11, 1853, 1754, 840
7SL4 4.63, 1853, 1652, 902
7STJ 4.52, 1853, 1672, 886
7STK 4.32, 1853, 1726, 860
7STI 4.33, 1853, 1678, 855
6PYH 4.66, 1899, 1648, 705

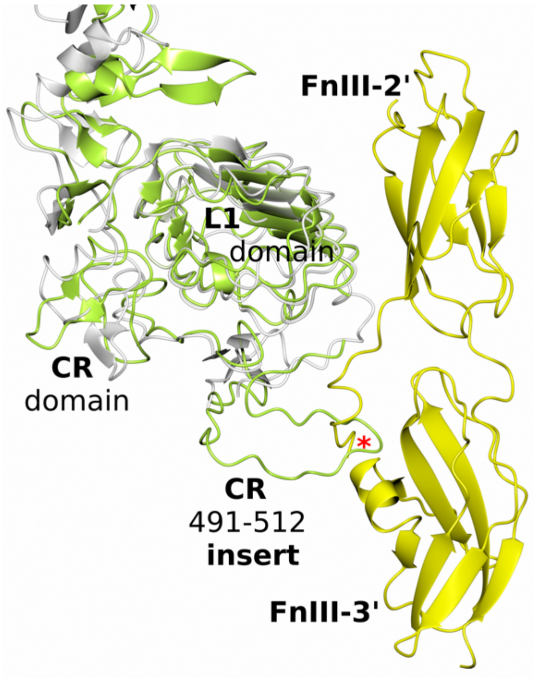

**Supplementary Information Figure 4. Putative steric hinderance caused by dmIR-specific CR 491-512 insert.**
The whole L1-CR domains of the dmIR-ECD were modelled here by AlphaFold[11], and the predicted (in green) and cryoEM structure L1 domains were superposed on their Cα atoms. Subsequently, the dmIR L1 was superposed on the L1 domain from the fully-down, insulin-free lower-arm of the hIR (PDB ID: 6HN5+6HN4) one insulin site 1 complex (in white) (targets as in Supplementary Note 1). This may suggest that the CR-insert in an unliganded dmIR protomer may cause a clash with FnIII-3' domain of the stem of the receptor (shown here in yellow, taken from the lower part of the hIR structure (PDB ID: 6HN4)). The red star indicates the overlapping regions.

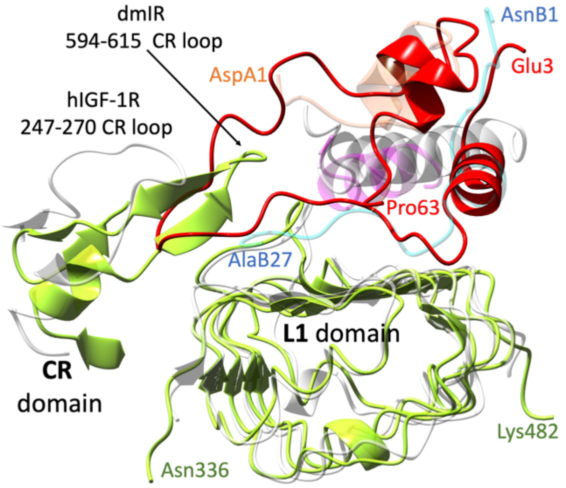

**Supplementary Information Figure 5. A putative clash of hIGF-1 C-domain with dmIR 594-615 CR loop**. dmIR in green, hIGF-1R in grey, hIGF-1 in red, DILP5 A- and B-chains in coral and blue, α–CT' in magenta. hIGF-1:hIGF-1R[3] (PDB ID: 7S0Q) complex structure was selected here due to a best definition of the loop region; superposition on the respective L1 domains (see Supplementary Information Note 1 for targets details).

**Supplementary Information Note 2. Comparison of some angular movements in dmIR-ECD and hIR.** Individual domains are depicted by boxes, Cα atoms selected as structural pivots are as black circles. The FnIII-1/2 domains are referred in the diagrams to as Fn1/2 for the brevity of the text.

***(A) L2-L1 domains angle:***

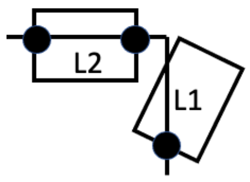

Cα pivots: 522TrpL1(183Trp) – 660GluL2(Glu329) – 792AspL2(456Asp) (hIR in brackets)

dmIR-ECD:
165.9° down Λ arm (B chain)
108.0° up Γ arm (A chain)

6SOF both T arms:
A chain 110.1°
B chain 113.1°

6HN5:

| | |
|---|---|
| E chain | 114.0° up Γ arm |
| D/F chain | 158.8° down Λ arm |

***(B) L2-Fn1-Fn2 for up Γ arms:***

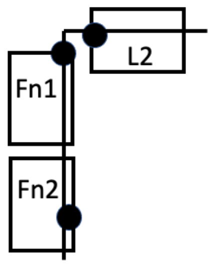

Cα pivots: 696Ala(360Leu) - 908Ile(Thr571) – 938Thr(Val604)

| | | |
|---|---|---|
| dmIR | A chain | 101.9° |
| 6SOF | C chain | 86.5° |
| | A chain | 90.0° |
| 6HN5 | C chain | 92.2° |
| 7PG0 | A chain | 99.0° |
| 7STH | A chain | 87.7° |

| | | |
|---|---|---|
| | B chain | 87.9° |
| 7STJ | B chain | 94.0° |

*(C) L1-Fn1-Fn2 for down Λ arms:*

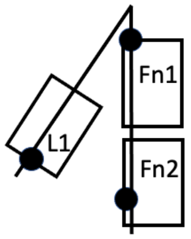

Cα pivots: 432LeuL1(104Leu) – 908IleF1(571Thr) – 938ThrF2(604Val); amino acids pivots in the L1 domain were selected for their structural similarity.

| | | | |
|---|---|---|---|
| dmIR | A chain | 60.4° | |
| 6HN5 | | 22.0° | L1 D chain, Fn1 F chain, Fn2 D chain |
| 7PG0 | B chain | 65.0° | |
| 7STJ | A chain | 50.9° | |
| 7STK | A chain | 53.4° | |
| 7PG3 | B chain | 35.4° | |
| 7PG4 | B chain | 72.4° | |

**Supplementary Information Note 3. Summary of the observed tertiary and quaternary trends among dmIR-ECD:DILP5 complex and h(m)IRs.**

The relative angles between L1-L2 domains, as determined by 522TrpL1(183Trp) – 660GluL2(Glu329) – 792AspL2(456Asp) angle, follow similar trends in all IRs. In dmIR they are 166° for the down Λ arm and 108.0° for the up *Γ* arm. In symmetrical T-form hIR, for example in the 6SOF structure, they are 110°/113° , while in the asymmetric one-insulin hIR complex (PDB ID: 6HN5+6HN4) they are 158.8° in the down Λ arm and 114.0° in the up *Γ* arm. The angles that depicts the 'rectangularity' between the FnIII-domains formed stem of the IRs (FnIII-1-FnIII-2-FnIII-3) and the L2 domain (represented by 695AlaL2(360Leu) – 908IleF1(571Thr) – 938ThrF2(604Val) angle) are within ca. 86° (6SOF) to 99° (7PG0) range in the hIRs, with 102° of the dmIR-ECD in the upper *Γ* arms following this trend.

The largest angular diversity is observed for the L1-FnIII-1-FnIII-2 angle for the down Λ dynamic protomers' arms, which reflects the range of the detachment of their L1 domain from the stem of the IR. As expected, this angle is lowest - 22° - for the fully IR-stem attached L1 domain in one-insulin at the upper site 1 complex (6HN5), attaining subsequently a broad range of values from 35° (7PG3)(3ins/IR), 65° (7PG0)(model 1 – 3 ins/IR), 72° (7PG4)(model 4 with two ins/hIR), and 51° and 53° in the hIR structures with site 1 down Λ-like arm insulin. Therefore, the 60° angle of the Λ arm in the DILP5:dmIR-ECD complex falls well within this range. The main – dmIR-specific - 'outlier' here is probably the furthest deviation of the L1-CR-L2 arm from the 'T'-plane of the IR, if measured, for example, between Arg 411(83) from the edge of the L2 as a reference points. Arg411 of the down-arm L2 domain is separated from its human equivalents by ~9-15 Å (after global SSM superpositions of the receptors; otherwise all above geometry indicators are based on specific superpositions targets of the appropriate domains.

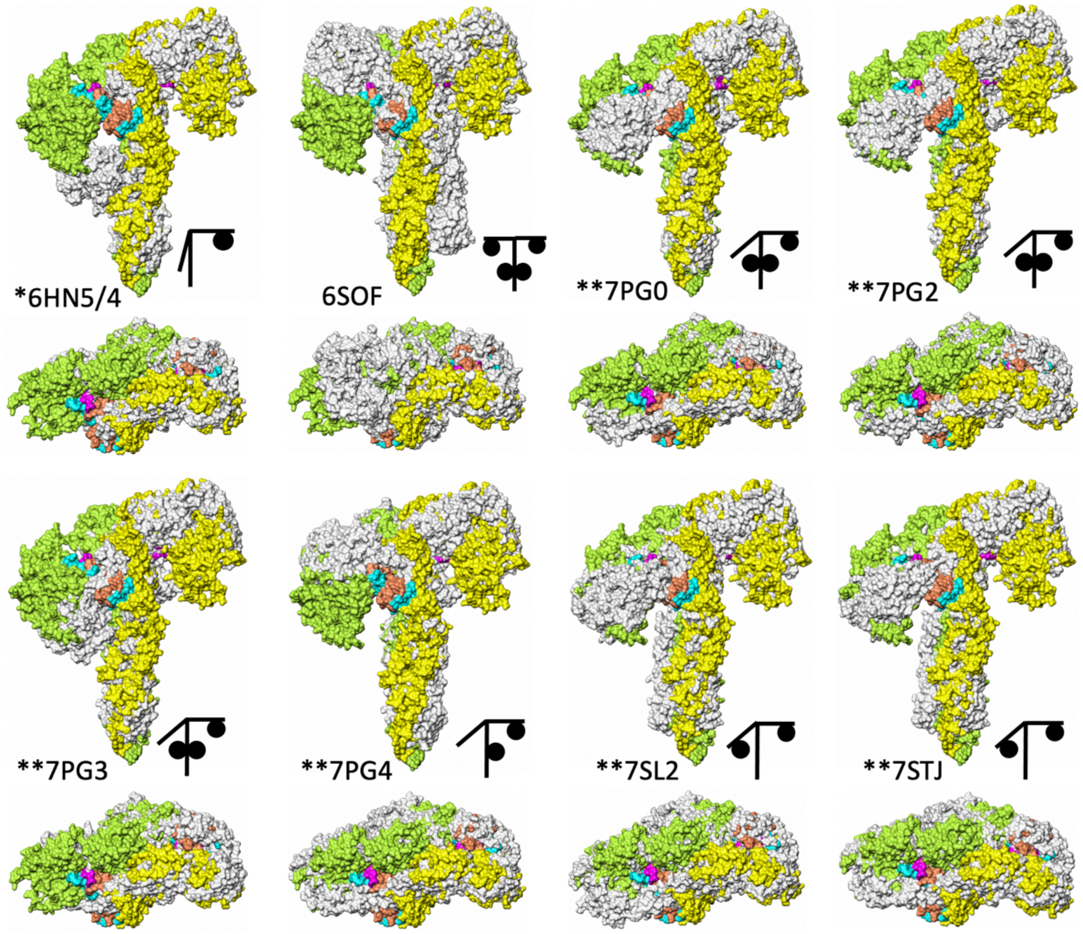

**Supplementary Information Figure 6. Global SSM superposition of dmIR-ECD and selected representative h/mIR structures** (including hormones); dmIR dynamic protomer in green, static protomer in yellow, h/mIR in white, DILP5's B-chains in blue, A-chains in coral, α-CT segment in magenta (rms details in Supplementary Information Note 1(E)). Stars indicate the source of the ECD; * - hIR-ECD with Leu-zippers, **-ECDs from full length h(m)IRs.

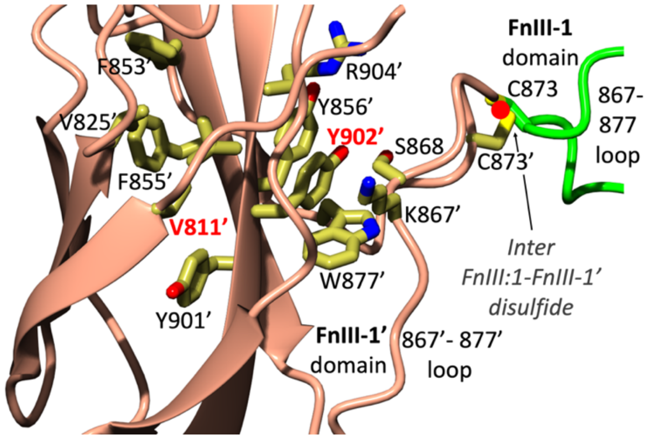

**Supplementary Information Figure 7. The possible impact of Y902C and V811D mutations[12] in on the dmIR.**
The FnIII-1' and FnIII-1 domains folds are in coral and green, respectively; red dot indicates the C873-C873' inter-protomers disulphide that crosslinks the FnIII-1 domains. Only the side chains from the hydrophobic cavities that surround Y982 and V811 are indicated.

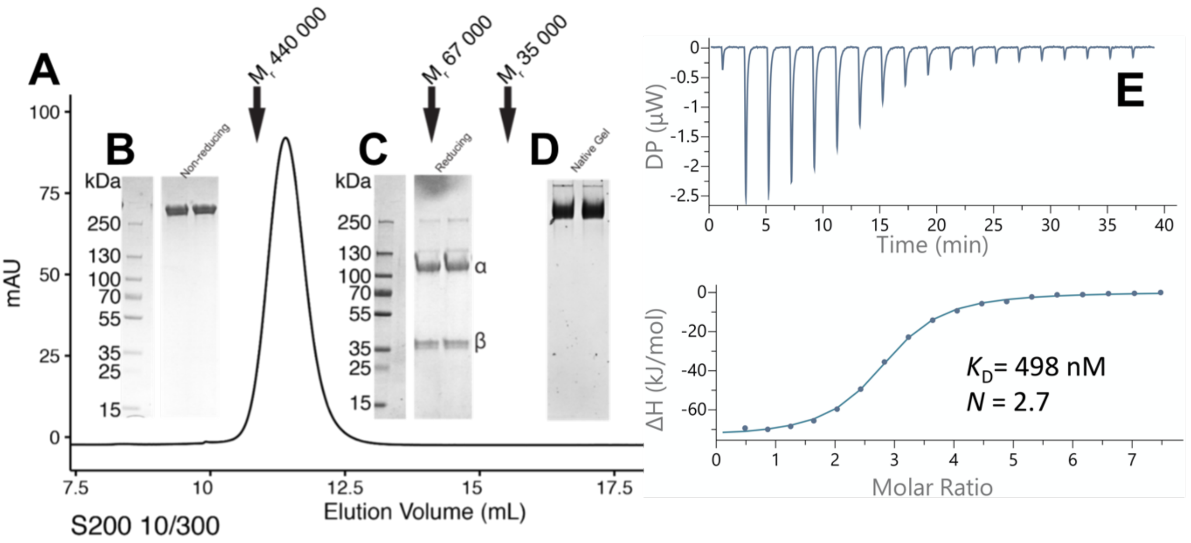

**Supplementary Information Figure 8. Purification and size exclusion chromatography of dmIR-ECD**. (A) Freshly purified dmIR ecto domain analyzed by SEC in 50 mM Tris, 150 mM NaCl, pH 7.4 using a Superdex S200 10/300 GL column (Cytiva). Arrows indicate elution volume of markers 440 000 Da (Ferritin), 67 000 (BSA) and 35 000 β-Lactoglobulin). (B) 4-12% SDS-PAGE of SEC peak containing fractions under non-reducing conditions (C) under reducing conditions and (D) 10% Native gel of SEC peak containing fraction. (E) μITC trace showing titration of DILP-5 hormone into dmIR-ECD. Data fitted to the "One set of sites" binding model. Source data are provided as a Source Data file.

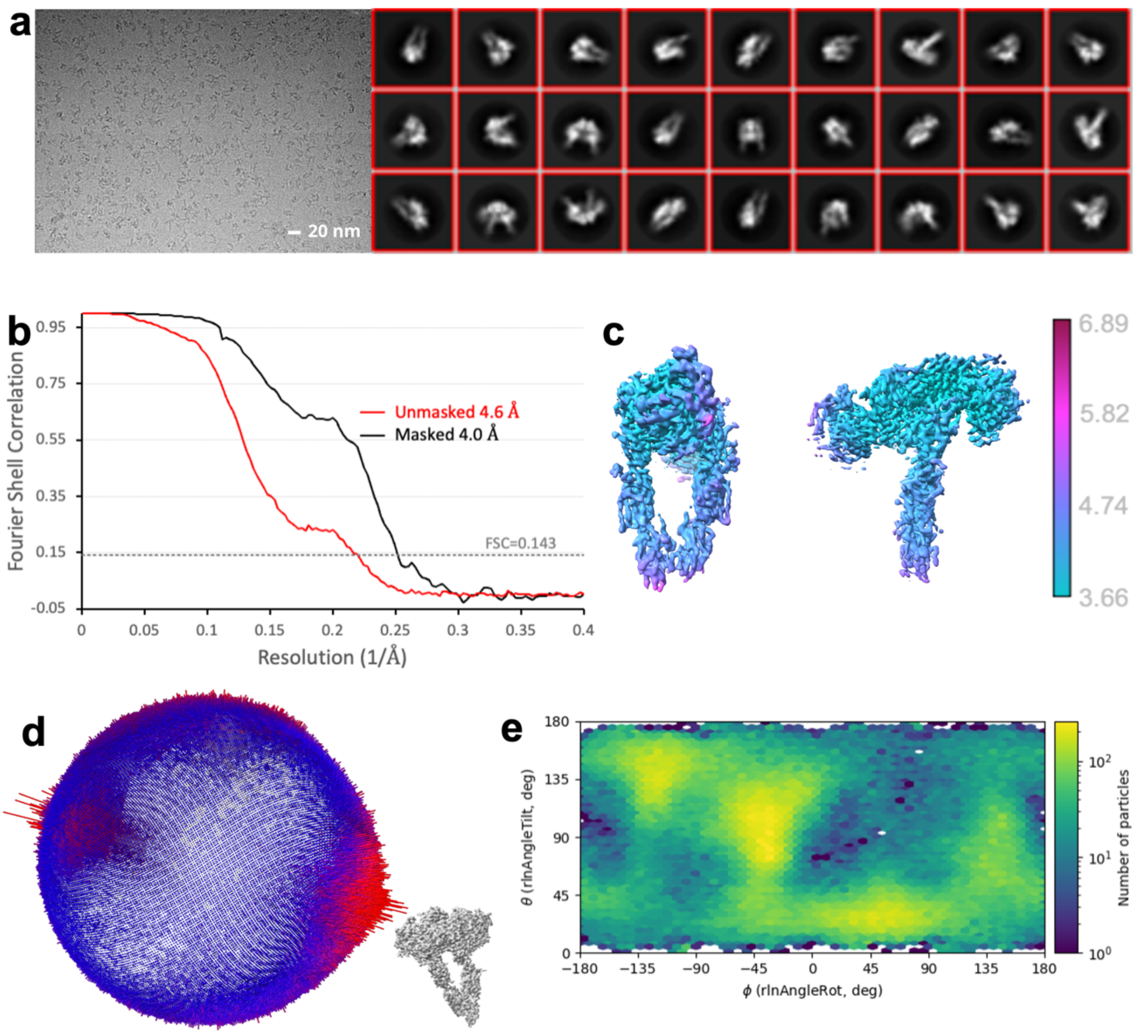

**Supplementary Information Figure 9**. **CryoEM acquisition and maps of dmIR-ecto:DILP5 complex.**

**a** Example micrograph of vitrified dmIR-ECD:DILP5 complex showing individual particles and some representative class averages on the right; the class associated with the apo-form of the dmIR-ECD is marked by a yellow circle. **b** Gold standard Fourier-Shell correlation resolution plot of masked and unmasked maps, with a cutoff at 0.143 indicated by a dashed line. **c** Map of dmIR-ECD:DILP5 complex with colour coded local resolution (right panel). **d** Distribution of particles views used in the final reconstruction projected on a sphere relative to the orientation shown on the lower right. **e** 2D histogram of Euler angles covered by a set of cryo-EM particles.

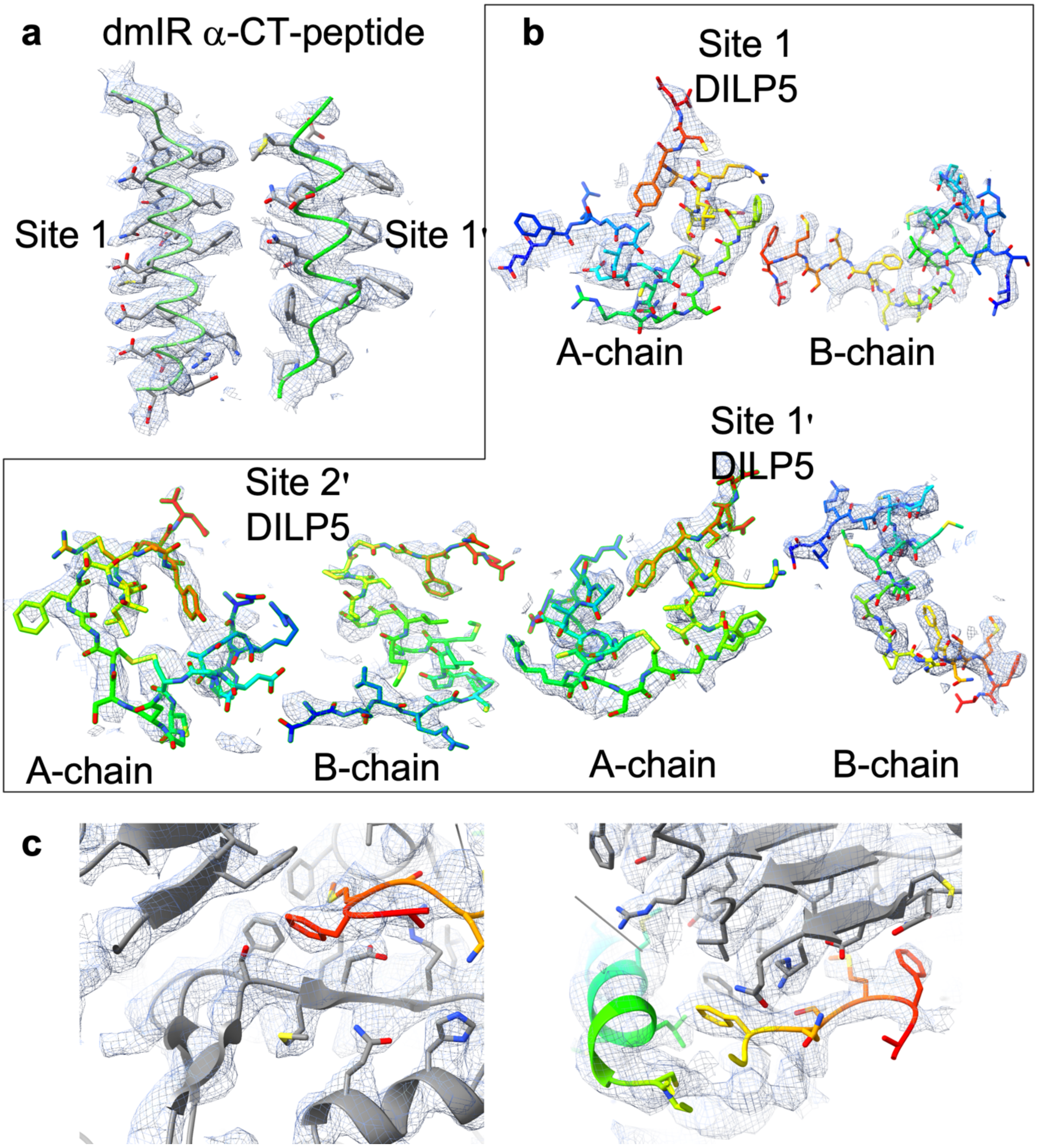

**Supplementary Information Figure 10.** Map and structural models for key dmIR-ECD:DILP5 complex structural elements and individual DILP5 hormone chains. **a** α-helical dIR α-subunit C-terminal region (α-CT) with cryo-EM map and model superposed. **b** Individual chains of each DILP5 hormone superposed on cryo-EM map at the same contour level. **c** Two views of residues from DILP5 Site 1 binding site in grey and hormone B-chain coloured in rainbow with cryo-EM map.

**Supplementary Information Table 1**. **Cryo-EM data collection, refinement and validation statistics**

| | dmIR-ECD:DILP5 complex (EMDB-16718) (PDB 8CLS) |
|---|---|
| **Data collection and processing** | |
| Magnification | - |
| Voltage (kV) | 300 |
| Electron exposure (e–/Å$^2$) | 0.829 (binned by 2) |
| Defocus range (μm) | 0.5 to 4 |
| Pixel size (Å) | 1.03 |
| Symmetry imposed | None |
| Initial particle images (no.) | 1.2m |
| Final particle images (no.) | 129,444 |
| Map resolution (Å) | 4.0 (0.143) |
| FSC threshold | |
| Map resolution range (Å) | 3.6 to 6.89 |
| **Refinement** | |
| Initial model used (PDB code) | AlphaFold 2.1, 2WFV |
| Model resolution (Å) | n/a |
| FSC threshold | n/a |
| Model resolution range (Å) | n/a |
| Map sharpening *B* factor (Å$^2$) | -50 |
| Model composition | |
| Non-hydrogen atoms | 28095 |
| Protein residues | 1840 |
| Ligands | 0 |
| *B* factors (Å$^2$) | |
| Protein | 45.76 |
| Ligand | n/a |
| r.m.s. deviations | |
| Bond lengths (Å) | 0.007 |
| Bond angles (°) | 0.829 |
| Validation | |
| MolProbity score | 1.91 |
| Clashscore | 4.13 |
| Poor rotamers (%) | 1.94 |
| Ramachandran plot | |
| Favored (%) | 91.97 |
| Allowed (%) | 8.06 |
| Disallowed (%) | 0.28 |

**Supplementary Information References**

1. Emsley, P., Lohkamp, B., Scott, W. G. & Cowtan K. Features and development of Coot. *Acta Crystallogr.* **D66**, 486-501 (2010).

2. Li, J., Choi, E., Yu, H. & Bai X-c. Structural basis of the activation of type 1 insulin-like growth factor receptor. *Nat. Commun.* **10**, 4567 (2019).

3. Xu, Y., Margetts, M. B., Venugopal, H., Menting, J. G., Kirk, N. S., Croll, T. I., Delaine, C., Forbes, B. E. & Lawrence, M. C. How insulin-like growth factor I binds to a hybrid insulin receptor type 1 insulin-like growth factor receptor. *Structure* **30**, 1-11 (2022).

4. Weis, F., Menting, J. G., Margetts, M. B., Chan, S. J., Xu, Y., Tennagels, N., Wohlhart, P., Langer, T., Müller, C.W., Dreyer, M. K. & Lawrence, M. C. The signalling conformation of the insulin receptor ectodomain. *Nat. Commun.* **9**, 4420 (2018).

5. Gutmann, T., Schäfer, I. B., Poojari, C., Brankatsch, B., Vattulainen, I, Strauss, M. & Coskün Ü. Cryo-EM structure of the complete and ligand saturated insulin receptor ectodomain. *J. Cell Biol.* **219**, e201907z10 (2020).

6. Uchikawa, E., Choi, E., Shang, G., Yu, H. & Bai, X.-c. Activation mechanism of the insulin receptor revealed by cryo-EM structure of the fully liganded receptor-ligand complex. *eLife* **8**, e48630 (2019).

7. Nielsen, J., Brandt, J., Boesen, T., Hummelshoj, T., Slaaby, R., Schluckebier, G. & Nissen, P. Structural investigations of full-length insulin receptor dynamics and signalling. *J. Mol. Biol.* **434**, 167458 (2022).

8. Li, J., Park, J., Mayer, J. P., Webb, K. J., Uchikawa, E., Wu, J., Liu, S., Zhang, X., Stowell, M. H. B., Choi, E. & Bai, X-c. Synergistic activation of the insulin receptor via two distinct sites. *Nat. Struct. Mol. Biol.* **29**, 357-368 (2022).

9. Xiong, X., Blakeley, A., Kim, J. H., Menting, J. G., Schäfer, I. B., Schubert, H. L., Agrawal, R., Gutmann, T., Delaine, C., Zhang. Y. W., Artik, G. O., Merriman, A., Eckert, D., Lawrence, M. C., Coskün, Ü., Fisher, S. J., Forbes, B. E., Safavi-Hemami, H., Hill, C. P. & Chou, H.-C. Symmetric and asymmetric receptor conformation continuum induced by a new insulin. *Nat. Chem. Biol.* **18**, 511-519 (2022).

10. Wu, M. *et al.* Functionally selective signaling and broad metabolic benefits by novel insulin receptor partial agonists. *Nat. Commun.* **13**, 942 (2022).

11. Jumper, J. *et al*. Highly accurate protein structure prediction with AlphaFold. *Nature* **596**, 583-589 (2021).

12. Banzai, K. & Nishimura, T. Isolation of a novel missense mutation in insulin receptor as a spontaneous revertant in ImpL2 mutants in *Drosophila*. *Development* **150**, dev201248 (2023).
